# Predator odor stress produces sex- and subpopulation-specific increases in alcohol drinking, anxiety-like behavior, and lateral hypothalamic *crh* expression

**DOI:** 10.1101/2025.03.20.644324

**Authors:** SM Bonauto, OR Brunke, FM Vassoler, MM Weera

## Abstract

Traumatic stress leads to maladaptive avoidance behaviors and alcohol misuse in some people. In rats, predator odor (“traumatic”) stress produces persistent avoidance of stress-paired contexts and escalated alcohol self-administration in some animals (Avoiders), but not others (Non-Avoiders). This mirrors the individual differences in stress responsivity and alcohol misuse seen in humans. Here, we used a quinine-adulterated alcohol drinking procedure to model compulsive-like alcohol drinking in humans. Male and female Wistar rats were given 12 weeks of intermittent access to 20% (v/v) alcohol, followed by three weeks of limited access. Rats were then indexed for avoidance using predator odor stress exposure, and limited access drinking resumed for three additional weeks after stress. During this period, the alcohol solution was adulterated twice weekly with increasing concentrations of quinine. More Avoidant males were more resistant to quinine adulteration and Avoider males increased in non-quinine alcohol drinking. Using ultrasonic vocalizations (USVs) as a measure of affective state, we found that Non-Avoider males emitted more lower frequency USVs (<32 kHz) preceding, during, and following predator odor stress. Finally, quantification of *crh, crhr1, crhr2, crhbp* gene expression in the lateral hypothalamus revealed a strong positive correlation between greater *crh* transcripts and avoidance in males and a positive correlation between *crh* transcripts and less anxiety-like behaviors in females. Together, these results suggest that the intersection of stress and compulsive-like alcohol drinking is sex-specific and dependent on individual differences in stress outcomes. This work reinforces the importance of considering sex differences in stress and alcohol use disorder research.

**Highlights:**

- Male Avoider rats show elevated two-bottle choice alcohol drinking after predator odor stress
- More avoidant males show more aversion-resistant alcohol drinking
- Female Avoider rats show heightened anxiety-like behavior 4 weeks after stress
- Low frequency USVs predict Non-Avoider behavior in male rats
- *Crh* expression in the LH is correlated with avoidance and alcohol drinking in male rats

## 1. Introduction

Living through a traumatic event leads to post-traumatic stress disorder (PTSD) in some individuals but not others (Goldstein et al., 2016). Key symptoms of PTSD include avoidance of trauma-related stimuli and increased negative affect after trauma (American Psychiatric Association, 2013). Individuals with PTSD are more likely to develop alcohol use disorder (AUD) (Straus et al., 2018), with about one-third of people diagnosed with PTSD at some point in their lives also meeting criteria for AUD (Blanco et al., 2013). Further, studies show that individuals with greater PTSD symptom severity consume more alcohol and are more likely to report that they drink to cope with negative symptoms (Lehavot et al., 2014; Ranney et al., 2021). While PTSD rates are higher in women after traumatic events (Goldstein et al., 2016). rates of AUD are generally greater in men (Erol & Karpyak, 2015), but this gap appears to be narrowing (Guinle & Sinha, 2020). The prevalence of comorbid AUD and PTSD across sex is nuanced, warranting further study (Gilpin & Weiner, 2017; Saraiya et al., 2023). Regardless, consensus falls on the idea that individuals with PTSD are particularly vulnerable to AUD due to stress-induced alterations in affective neural circuitry that increase negative affect and drive alcohol drinking via negative reinforcement (Koob, 2013a, 2013b, 2014; Kwako, Schwandt, et al., 2015; Straus et al., 2018). Animal models are necessary to explore the neurobiology of these disorders with consideration of potential sex differences.

In lab rats, some aspects of PTSD symptomatology and individual differences in stress responsivity are recapitulated via a predator odor stress model. In this paradigm, a single, inescapable exposure to bobcat urine produces persistent avoidance of the odor-paired context and prolonged increases in anxiety-like behavior in a subset of outbred Wistar rats, termed “Avoiders” (Edwards et al., 2013; Whitaker & Gilpin, 2015; Schreiber et al., 2017; Albrechet-Souza & Gilpin, 2019; Albrechet-Souza et al., 2020; Weera et al., 2023). All predator odor-exposed animals show evidence of acute stress, such as elevated serum corticosterone and ACTH in the hours after predator odor exposure, blunted body weight gain, and heightened anxiety-like behavior over a period of several days after stress (Whitaker & Gilpin, 2015; Weera et al., 2020). However, only Avoiders show escalated alcohol self-administration (SA) (Edwards et al., 2013; Whitaker & Gilpin, 2015; Schreiber et al., 2017; Weera et al., 2020, 2023) and hyperalgesia after stress (Itoga et al., 2016). The heightened anxiety-like behavior after stress also appears to be more persistent in Avoider rats, lasting at least up to 9 days post-stress (Weera et al., 2023). Studies also show that Avoider or Non-Avoider status remains stable over weeks after stress and is not altered by repeated odor exposure (Schreiber et al., 2017; Weera et al., 2020). Importantly, we highlight that most of this work was performed in male rats, although more recent studies point towards similar behavioral phenotypes in female rats following predator odor stress (Albrechet-Souza et al., 2020; Weera et al., 2023).

Neurocircuit studies using this animal model implicate the lateral hypothalamus (LH) as an important node downstream of the amygdala for mediating Avoider-related stress responses. For instance, activity of inputs into the LH from the central amygdala (CeA) is required for supporting avoidance of a stress-paired context (Weera et al., 2021), and inhibition of CeA-to-LH inputs that express corticotropin-releasing factor (CRF) type-1 receptors (CRF1) is sufficient for normalizing the escalated alcohol self-administration and anxiety-like behavior in male and female Avoider rats. Within the LH, antagonism of CRF1 reduces binge-like alcohol drinking in male mice (Bendrath et al., 2025) and rescues heightened anxiety-like behavior after acute stress (Eghtesad et al., 2022). Together, these findings implicate the CRF neuropeptide system within the LH in driving post-stress alcohol drinking and anxiety-related behaviors.

The main goal of this study was to test the hypothesis that both male and female Avoider rats will show increased free-choice alcohol drinking after stress. In addition to free-choice alcohol drinking, we also tested Avoider, Non-Avoider, and unstressed Control rats’ consumption of alcohol solutions that were adulterated with an aversive stimulus (i.e., quinine). Humans with AUD typically show sustained alcohol consumption despite negative consequences (also called ‘compulsive-like’ alcohol drinking) (Sinha, 2009; Koob & Volkow, 2010; Vendruscolo et al., 2012; American Psychiatric Association, 2013), and the quinine-adulterated alcohol drinking procedure is a model of aversion-resistant drinking (Simms et al., 2008; De Oliveira Sergio et al., 2023). Given that alcohol drinking may be driven by negative emotional states, we tested the enduring nature of stress-induced anxiety-like behavior in Avoider and Non-Avoider rats, weeks after predator odor exposure, as well as their emotional states before stress, during stress, and during expression of avoidance or non-avoidance of stress-paired contexts using ultrasonic vocalizations (USVs) as a proxy. Finally, we quantified levels of *crh, crhr1, crhr2,* and *crhbp* in the LH. The findings here contribute to the growing understanding of subpopulation- and sex-specific effects of stress.

## 2. Methods

### 2.1 Animals

Adult (8 weeks old) male and female Wistar rats (Charles River, Kingston, NY) were single-housed in a temperature- and humidity-controlled vivarium (22°C) on a 12-hour reversed dark/light cycle (lights off at 9 am EST). Rats were provided with tap water and chow (Teklad 2919) *ad libitum.* All rats were handled and weighed at least once each week during the duration of the experiments. Behavioral tests occurred during the dark period. All experiments were approved by the Tufts University Institutional Animal Care and Use Committee (IACUC).

### 2.2 Predator Odor Conditioned Place Aversion

All rats underwent a 4-day predator odor conditioned place aversion (CPA) procedure as previously described (Edwards et al., 2013; Weera et al., 2023). Briefly, on the first day, rats explored a three-chamber arena with a center zone and two chambers with distinct tactile (mesh vs. holes) and visual (stripes vs. dots) stimuli for 5 minutes in a dimly lit room (maximum light 13 lumens; Pretest). Duration spent in each chamber was recorded by experimenters blinded to rat identities. Placement of three paws inside of striped or dotted chambers was required for an entry to be counted. On the second day, individual rats were confined to either the stripes or dots chamber for 15 minutes (neutral chamber). On the third day, individual rats were confined to the other chamber with a bobcat urine-soaked paper towel (or nothing for unstressed controls) for 15 minutes (odor chamber). On the final day, rats were again allowed to explore the entire arena for 5 minutes (Posttest). The change in duration spent in the odor-paired chamber from Pretest to Posttest was recorded for odor-exposed (stressed) rats. Stressed rats with a decrease in time spent in the odor-paired chamber that was greater than 10 seconds were classified as “Avoiders.” While other stressed rats were designated “Non-Avoiders” (Edwards et al., 2013; Albrechet-Souza & Gilpin, 2019). Non-odor exposed animals were considered unstressed Controls. The behavioral apparatus was cleaned with chlorine dioxide between each animal on all days, and with soap, water, and an enzyme-based odor remover (PureAyre Odor Eliminator, Clean Earth Inc., Kent, WA) at the end of predator odor exposure.

### 2.3 Ultrasonic Vocalization Recording and Analysis

During all days of predator odor place conditioning, ultrasonic vocalizations (USVs) were recorded from the actively behaving rat. An Avisoft Bioacoustics Condenser Microphone (CM16/CMPA, Avisoft Bioacoustics UltraSoundGate Model 116Hb) was suspended 18 inches above the center of the place conditioning apparatus and Avisoft RECORDER software was used to monitor recording. Audio files were processed in DeepSqueak using the built-in Long Rat and Rat neural networks to generate call detection files which were manually edited for vocalization identification accuracy by a team of analysts blinded to rat condition (Coffey et al., 2019). Data files were analyzed for quantity of calls as well as number of calls with the average principal frequency of less than 32 kHz which is the maximum range for vocalizations typically emitted in aversive situations (Brudzynski, 2001; Burgdorf et al., 2008; Takahashi et al., 2010). For the 15-minute odor day, only the first 5 minutes of the audio file was analyzed to allow comparison to the 5-minute pre- and post-stress sessions.

### 2.4 Alcohol Drinking

Procedures were adapted from published studies (Simms et al., 2008; De Oliveira Sergio et al., 2023). Sixty rats (30 male, 30 female) were single housed to accurately measure free-choice water and alcohol consumption in each rat. Rats were given 12 weeks of Intermittent Access two-bottle choice alcohol (20% v/v) drinking on a Monday, Wednesday, Friday schedule, followed by 3 weeks of Limited Access (20-minute) two-bottle choice drinking on a Monday to Friday schedule. On the third week of Limited Access drinking, the alcohol solution was adulterated with 10 mg/L quinine monohydrochloride dihydrate (Sigma-Aldrich) on Tuesday and Thursday. Next, rats were counterbalanced into Stress and unstressed Control groups based on average consumption of quinine adulterated alcohol (i.e., Quinine Baseline) and subjected to the 4-day (Monday-Thursday) predator odor CPA procedure described above (2.2). Limited Access alcohol drinking resumed on the next day (Friday) and for three additional weeks on a Monday-Friday schedule. Across these 3 weeks, the alcohol solution was adulterated with increasing concentrations of quinine (10 mg/L, 60 mg/L, 100 mg/L) twice each week (Figure 1A).

**Figure 1.**
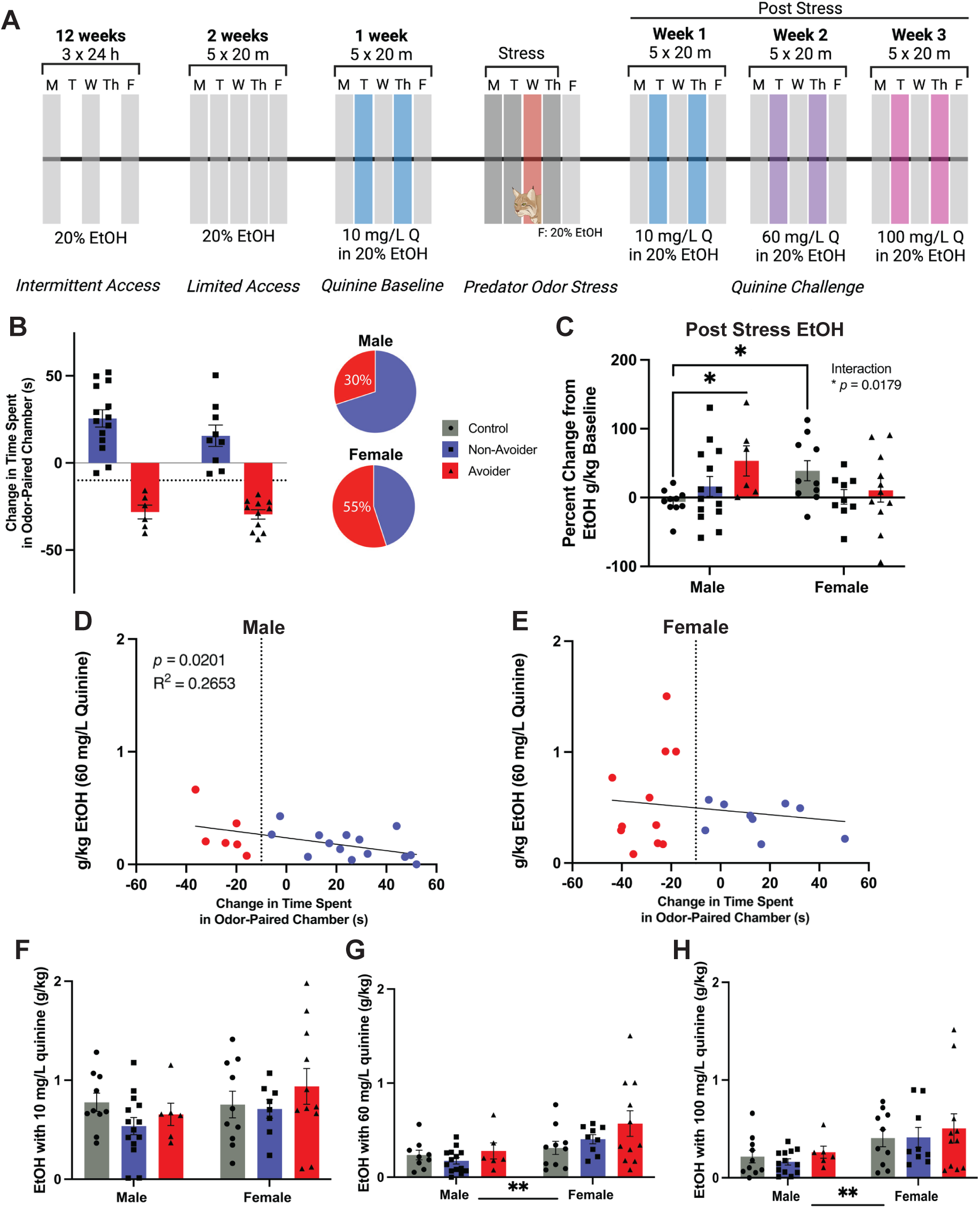
Avoidant males show increased free-choice alcohol drinking and quinine-resistant drinking. **A)** Timeline schematic of the experiment. **B)** Distribution of avoidance vs. non-avoidance of predator odor stress-paired contexts (male Avoiders = 6, female Avoiders = 11). **C)** Post-stress alcohol consumption averaged across 3 weeks of post-stress non-quinine drinking sessions. Expressed as percent change from pre-stress alcohol-only drinking sessions during the Quinine Baseline week (i.e., averaged across Monday, Wednesday, and Friday of that week). **D)** Relationship between change in time spent in odor-paired chamber (avoidance score) and consumption of alcohol adulterated with 60 mg/L in male and **E)** female rats. **F)** Post-stress consumption of alcohol adulterated with 10 mg/L, **G)** 60 mg/L, or **H)** 100 mg/L quinine. Error bars represent Mean ± SEM. \**p* <0.05, \*\**p* < 0.01. Created in BioRender. Bonauto, S. (2025) https://BioRender.com/w93w369.

### 2.5 Behavioral Assays Following Alcohol Drinking

Rats were subjected to a 5-minute elevated plus maze (EPM) test in a dim (maximum 40 lumens in open arms), novel room 5-6 days after the final drinking session. Rats were introduced to the center zone of the EPM and allowed to explore undisturbed. Behavior was recorded by an overhead video camera and processed using Ethovision XT (Noldus Information Technology, v17.5) to score time in open arms, entries into each zone, and locomotion. Hand scoring for stretch attempts was conducted by an independent experimenter blinded to rat condition. Criteria for stretch attempts included entry of the front two paws into the open arms followed by retreat.

### 2.6 Quantitative PCR (qPCR)

Four weeks following predator odor stress, rats were sacrificed, and brains were snap frozen in 2-methyl butane and stored at -80°C until dissection. Bilateral punches (1 mm diameter, 0.5mm depth) of the lateral hypothalamus (LH) were collected at +2.2 A/P, ±2.0 M/L, -8.8 D/V from Bregma according to the atlas of Paxinos and Watson. for Quantitative PCR (qPCR) across all rats and stored at -80°C. RNA was extracted using the Qiagen RNeasy RNA extraction kit (Hilden, Germany) according to manufacturer’s instructions.

<u>RTqPCR:</u> Complementary DNA (cDNA) was synthesized using the RETROscript kit from Applied Biosystems (Carlsbad, CA, USA). PCR was performed using a Quant3 (Fisher Scientific) under standard amplification conditions: 2 min at 50 °C, 10 min at 95 °C, 40 cycles of 15 s at 95 °C and 60 s at 60 °C. All PCR primers were TaqMan® Gene Expression Assays purchased from Applied Biosystems. The amplification efficiency of each of these assays has been validated by Applied Biosystems and averages 100% (± 10). Assay ID and accession numbers were as follows: *gapdh:* Rn01775763_g1*; crh:* Rn01462137_m1*; crhbp:* Rn00594854_m1*; crhr1:* Rn00578611_m1; *crhr2:* Rn00575617_m1. Final quantification of mRNA was obtained using the comparative cycle threshold (CT) method (User Bulletin #2, Applied Biosystems). Data are reported as fold change in transcription relative to a calibrator cDNA. In brief, the housekeeping gene (*gapdh*) was used as an internal control against which each target signal was normalized (ΔCT). Validation studies confirmed that the raw CT values of *gapdh* did not vary by treatment group (Supplemental Figure S3A). The ΔCT was then normalized against a calibrator (i.e. the mean of the control male group for the target gene) to provide the ΔΔCT relative to the control group. Finally, data are transformed 2^ ΔΔCT and expressed as fold change.

### 2.7 Statistical Analysis

All data were analyzed using omnibus ANOVAs with Sex and Stress Group as between-subjects factors, and Day as a within-subjects factor for alcohol drinking. Significant main effects were followed up with lower order ANOVAs and Tukey’s post hoc tests. Correlations were performed using simple linear regression analyses. Alpha was set at 0.05. Outliers were determined using Grubbs Outlier Test (≤1/group). Analysis was performed in Prism 10 (GraphPad) and SPSS Statistics (IBM). Data are shown as Mean ± SEM.

## 3. Results

### 3.1 Male Avoider rats show elevated free-choice alcohol drinking

All rats had two-bottle choice intermittent access to 20% (v/v) alcohol in the home cage for 12 weeks (Intermittent Access) followed by two weeks of Limited Access and one week of Quinine Baseline (Figure 1A). Alcohol intake during Intermittent Access, 24-hour drinking sessions was similar between sexes and between animals that were subsequently counterbalanced into Stress or unstressed Control groups (Supplemental Figure S1A). In the Limited Access phase in which rats had alcohol access for 20 min 5x/week, alcohol intake increased over two weeks (Repeated Measures Day x Sex ANOVA, significant main effect of Day, F (4.193, 243.2) = 10.09, *p*<0.0001) and stabilized (Supplemental Figure S1B). A mixed-effects analysis (Sex x Day) of percent preference for alcohol (volume of alcohol consumed/total liquid consumed*100) during Limited Access revealed that females had greater preference for alcohol (F (1, 58) = 4.220, *p=*0.0445) and, for both sexes, alcohol preference increased across drinking sessions (F (4.166, 239.8) = 4.671, *p*=0.0010) (Supplemental Figure S1C). A mixed-effects analysis was utilized to correct for five randomly missing values due to water bottle leakage.

Rats were assigned to Control or Stress conditions in a counterbalanced fashion based on their average consumption of quinine-adulterated alcohol the week before stress (Quinine Baseline; Supplemental Figure S1D). Following predator odor stress, 30% of male rats and 55% of female rats showed avoidance of stress-paired contexts (Figure 1B). A subset of animals was kept as unstressed, handling Controls and were tested in parallel to stressed animals.

One purpose of this experiment was to test the hypothesis that Avoiders would show increased free-choice unadulterated alcohol drinking. Post-stress changes in unadulterated alcohol drinking were quantified as a percent change relative to the pre-stress Alcohol Baseline (i.e., average alcohol consumed over the Monday, Wednesday, and Friday before the week of stress procedures). Change from Alcohol Baseline was averaged for Monday, Wednesday, Friday for each post-stress week. Outliers were excluded and then the post-stress unadulterated alcohol drinking values for each week were averaged. We found that male Avoider rats drank more alcohol (without quinine adulteration) across 3 weeks following predator odor stress compared to unstressed Controls (Stress Group x Sex ANOVA, interaction, F (2,54) = 4.341, *p* = 0.0179; Tukey’s post-hoc, *p*=0.0407 for Avoiders vs. Controls) (Figure 1C). Alcohol intake across Monday-Wednesday-Friday of each post-stress week remained stable (i.e., no statistical effect of day or week).

Another purpose of this experiment was to test the hypothesis that Avoider rats would show greater compulsive-like alcohol drinking using a free-choice quinine-adulterated alcohol drinking procedure. Following the week of predator odor stress CPA, rats underwent three weeks of Monday to Friday limited access drinking sessions, with the alcohol solution adulterated with quinine twice weekly on Tuesdays and Thursdays. The concentration of quinine increased from 10 mg/L (Week 1 post-stress) to 60 mg/L (Week 2 post-stress) to 100 mg/L (Week 3 post-stress) (Figure 1A). Within each sex, within subjects RM two-way ANOVAs for intake of quinine-adulterated alcohol across all quinine concentrations (i.e., across all three weeks post-stress) showed significant main effects of quinine concentration (Stress Group x Quinine Concentration; males: F (1.407, 37.98) = 39.32, *p*<0.0001; females: F (1.560, 42.11) = 20.48, *p*<0.0001; Supplemental Figure S1E, F). Further analyses tested the effects of Stress Group on quinine-adulterated alcohol drinking within each quinine concentration.

In male rats (Figure 1D), we found an inverse correlation between avoidance scores and intake of quinine-adulterated alcohol, such that greater avoidance of stress-paired contexts predicted greater intake of alcohol that was adulterated with 60 mg/L quinine (R^2^=0.2653, *p=*0.0201). This relationship was not detected in female rats (R^2^=0.02424, *p*=0.5122; Figure 1E).

There were no stress group or sex differences in alcohol consumption when the alcohol solution was adulterated with a mild dose of quinine (10 mg/L) (Figure 1F). With moderate quinine challenge (60 mg/L), female rats consumed more alcohol than males (Stress Group x Sex ANOVA, main effect of Sex, F (1, 53) = 8.877, *p*=0.0044). (Figure 1G). At a higher quinine dose (100 mg/L), female rats still consumed more alcohol than males, but there were no Stress Group effects (Stress Group x Sex ANOVA, significant main effect of Sex, F (1, 53) = 8.371, *p=*0.0055) (Figure 1H).

### 3.2 Female Avoider rats show heightened anxiety-like behavior 4 weeks after predator odor stress

The purpose of this experiment was to test the enduring nature of stress-induced anxiety-like behavior in male and female Avoider vs. Non-Avoider rats. Animals were subjected to an EPM test 28 days after stress (Figure 2A). In general, female rats spent more time in the open areas (Stress Group x Sex ANOVA, main effect of Sex, F (1, 51) = 25.42, *p*<0.0001) (Figure 2B), and had more open arm entries than male rats (Stress Group x Sex ANOVA, main effect of Sex, F (1, 51) = 42.43, *p*<0.0001) (Figure 2C). Measurements of stretch attempts showed that female Avoider rats had more aborted attempted entries into the open arms of the plus maze (Stress Group x Sex ANOVA, interaction, F (2, 51) = 4.066, *p*=0.0230; Tukey’s post-hoc in female Avoiders vs. Controls, *p*=0.0198), indicating elevated anxiety-like behavior in these animals (Figure 2D). Additionally, both Avoider and Non-Avoider females executed more stretch attempts than males in the same stress groups (Tukey’s post-hoc Avoiders male vs female: *p* = 0.0005; Non-Avoiders male vs female: *p* = 0.0406). Finally, distance moved throughout the assay was similar across stress groups, though females showed more locomotion than males (Stress Group x Sex ANOVA, main effect of Sex, F (1, 50) = 27.32, *p*<0.0001) (Figure 2E).

**Figure 2.**
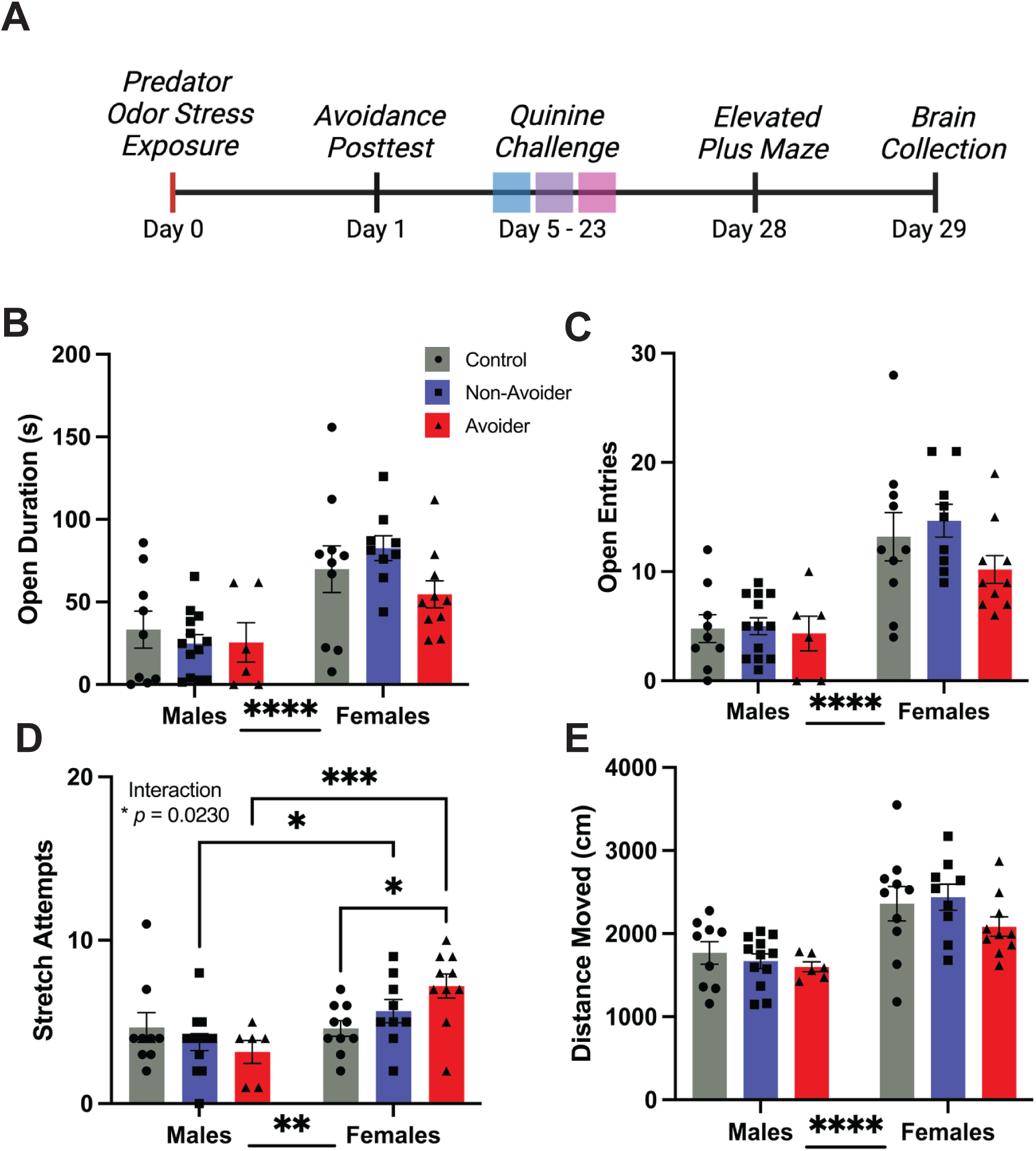
Female Avoider rats show heightened anxiety-like behavior 28 days after stress. **A)** Timeline schematic (extension of Figure 1A). **B)** Duration in open arms of the EPM. **C)** Number of entries into open arms of the EPM. **D)** Number of stretch attempts (front two paws without entry) into the open arms of the EPM. **E)** Distance moved during EPM. Error bars represent Mean +/- SEM. \**p* <0.05, \*\**p* < 0.01, \*\*\**p* < 0.001, \*\*\*\**p* < 0.0001. Created in BioRender. Bonauto, S. (2025) https://BioRender.com/v69x870.

### 3.3 Male Non-Avoider rats emit more <32 kHz ultrasonic vocalizations before, during, and after predator odor stress

The purpose of this experiment was to track USVs emitted by Avoider and Non-Avoider rats throughout predator odor place conditioning. During place conditioning, USVs were recorded and analyzed for principle frequency and total number of vocalizations. Throughout the place conditioning procedure, there were no group differences in the number of USVs emitted (Supplemental Figure S2A-C) and little change in the amount or type of USVs emitted by each individual rat from Pretest to Posttest (Supplemental Figure S2D, E). However, there were robust differences between Stress Groups and Sexes when we analyzed USVs within distinct frequency bands (Figure 3A). During the Pretest, males emitted significantly more <32 kHz USVs than females (Stress Group x Sex ANOVA, main effect of Sex, F (1, 51) = 9.264, *p*=0.0037; effect of Stress Group, F (2, 51) = 2.938, *p* = 0.0620). A direct comparison between the two subpopulations of stressed animals (i.e., Avoiders vs. Non-Avoiders) showed that males that would become Non-Avoiders with future odor stress emitted a higher percentage of <32 kHz vocalizations compared to Avoiders (unpaired t-test, t=2.383, df=16, *p*=0.0299) (Figure 3B). In males, there was also a positive correlation between eventual avoidance scores and percent of <32 kHz USVs during Pretest, such that a higher percentage of USVs in the <32 kHz range during Pretest predicted greater non-avoidance of a stress-paired context (R^2^=0.4371, *p*=0.0028; Figure 3C).

**Figure 3.**
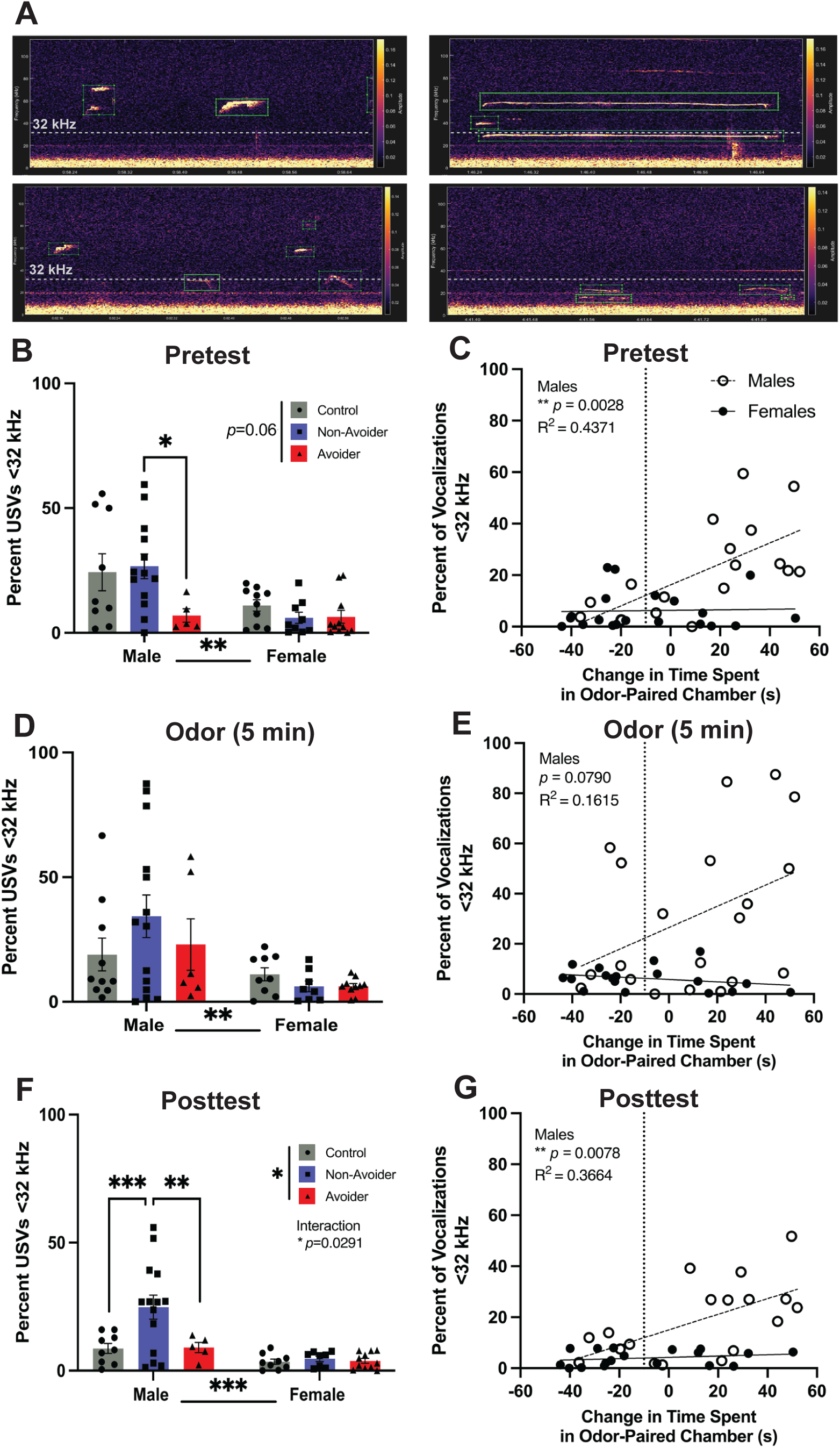
Non-Avoider males emit more <32 kHz USVs at all time points. **A)** Examples of <32 kHz USV spectrograms. **B)** Percentage of USVs with principle frequencies <32 kHz in the **B)** Pretest, **D)** first 5 minutes of odor exposure, **F)** Posttest. **C)** Relationship between change in time spent in odor-paired chamber (avoidance score) and percentage of vocalizations <32 kHz in the **C)** Pretest, **E)** first 5 minutes of odor exposure, **G)** Posttest. Error bars represent Mean +/- SEM. \**p* <0.05, \*\**p* < 0.01, \*\*\**p* < 0.001. Created in BioRender. Bonauto, S. (2025) https://BioRender.com/q64q502.

During predator odor exposure, vocalizations from the first 5 minutes of the recording were quantified. Males again emitted a greater percentage of <32 kHz vocalizations than females (Stress Group x Sex ANOVA, main effect of Sex, F (1, 51) = 9.919, *p*=0.0027), but vocalizations between stress groups were not significantly different (Figure 3D). Correlations between eventual avoidance scores and percent USVs <32kHz were also not significant (but in males, R^2^=1615, *p=*0.0790; Figure 3E).

During Posttest, 24 hours after predator odor stress, Non-Avoider males continued to emit a greater percentage of <32 kHz vocalizations (Stress Group x Sex ANOVA, interaction F (2, 50) = 3.800, *p*=0.0291, Tukey’s post hoc in male Control vs Non-Avoiders *p*=0.0007 and male Avoiders vs Non-Avoiders *p*=0.0073; Figure 3F). Additionally, there was a positive correlation between avoidance scores and <32 kHz USVs for males in the Posttest, such that less avoidant animals emitted a greater percentage of <32 kHz USVs (*p*=0.0078, R^2^=0.3664; Figure 3G).

### 3.4 Gene Expression Changes in the Lateral Hypothalamus

Genes associated with stress and the stress response were examined in the lateral hypothalamus 4 weeks after predator odor exposure. Levels of *crh* were significantly increased in male Avoiders compared with Non-Avoiders (*p*=0.025) and Controls (p=0.019) (Stress Group x Sex ANOVA, interaction, (F (2,51) = 5.654; *p*=0.0061; Figure 1A). Tukey’s post hoc multiple comparisons test also showed that female Control animals had higher *crh* compared to male Controls (*p*=0.016) and female Avoiders had less *crh* than male Avoiders (*p*=0.028). Expression of *crhbp* was also examined (Figure 4B). A two-way ANOVA revealed a significant main effect of Sex (F (1,47) = 4.305; *p*=0.0435).

**Figure 4.**
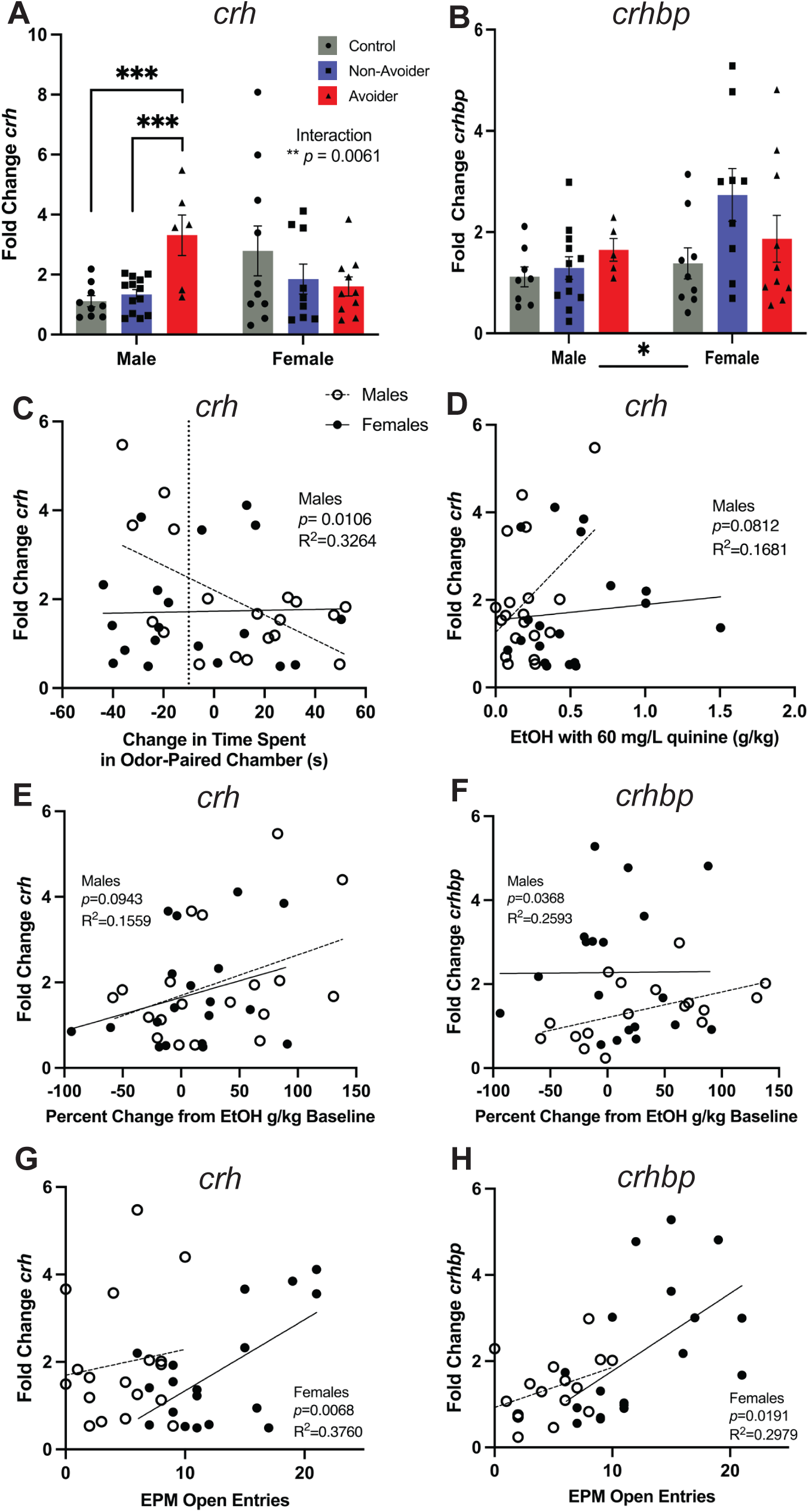
Gene expression changes in the lateral hypothalamus. **A)** Fold change in mRNA expression of *crh* and **B)** *crhbp*. **C)** Relationship between crh expression and avoidance score or **D)** amount of 60 mg/L quinine in alcohol consumed in post stress week 2. **E)** Correlation between post-stress change in unadulterated alcohol drinking relative to Alcohol Baseline and *crh* expression or **F)** *crhbp*. **G)** Relationship between open arm entries in the EPM and *crh* expression or **H)** *crhbp*. Error bars represent Mean +/- SEM. \**p* <0.05.

To test if there was a relationship between gene expression and post-stress behavioral changes, correlations were performed using all stressed rats. In male rats, Avoidance scores were negatively correlated with *crh* expression in the LH, such that greater avoidance scores predicted higher LH *crh* expression (R^2^=0.3264; *p*=0.0106; Figure 4C). In addition, the amount of quinine (60 mg/L) adulterated alcohol consumed tended to be positively correlated with *crh* expression in males (R^2^=0.1681; *p*=0.0812; Figure 4D). With regards to post-stress consumption of unadulterated alcohol in male rats, there was a weak relationship between percent change in alcohol drinking and *crh* expression (R^2^=0.1559; *p*=0.0943; Figure 4E), but the relationship between percent change in drinking and *crhbp* expression was significant (R^2^=0.2593, *p*=0.0368; Figure 5F). In females, there was a positive correlation between EPM open arm entries and *crh* expression (R^2^=0.3760, p=0.0068; Figure 4G) as well as with *crhbp* expression (R^2^=0.2979; p=0.0191; Figure 4H).

The expression levels of *crhr1* and *crhr2* in the LH were similar across Stress Groups and Sexes (Supplemental Figure S3B, C). However, in female stress-exposed rats, greater expression of *crhr1* in the LH predicted more open arm entries in the EPM (R^2^=0.2185, *p*=0.0505, Supplemental Figure S3D).

## Discussion

Individuals with PTSD are more susceptible to AUD, yet the mechanisms and neurobiological factors of this phenomenon are unclear (Goldstein et al., 2016; Straus et al., 2018). In the present study, we used a predator odor stress model that recapitulates some aspects of PTSD, such as persistent avoidance of stress-paired stimuli in the absence of the stressor (i.e., conditioned avoidance; Albrechet-Souza and Gilpin, 2019) to test the effects of an acute, potentially ‘traumatic,’ stressor on free-choice alcohol drinking; aversion-resistant alcohol drinking; anxiety-like behavior; ultrasonic vocalizations before stress, during stress, and during expression of conditioned avoidance; and CRF system gene expression in the lateral hypothalamus in male and female rats. We found that only subsets of male (30%) and female rats (55%) developed conditioned avoidance of a stress-paired context, reflecting individual differences in stress responsivity also seen in humans, and similar to previous findings in rats (e.g., Albrechet-Souza et al., 2020). Male Avoider rats showed elevated free-choice alcohol drinking and were more likely to show aversion-resistant alcohol drinking. Female Avoider rats showed signs of heightened anxiety-like behavior 4 weeks after odor exposure. Using USV recordings, we found that increased vocalizations in the <32 kHz range before stress and during testing for conditioned avoidance predicted more non-avoidance behavior in male but not female rats. In the LH, male Avoiders had elevated *crh* gene expression, whereas all female stressed and unstressed rats had more *crhbp*, when brains were collected 4 weeks after stress (or handling control procedures). Overall, these findings expand our understanding of Avoider vs. Non-Avoider phenotypes after stress, allowing us to embark on novel, mechanistic research avenues to elucidate the neurobiology of stress-alcohol interactions in subpopulations of animals.

Previous studies reliably demonstrate that Avoider rats show heightened alcohol operant self-administration after predator odor stress (Edwards et al., 2013; Schreiber et al., 2017; Weera et al., 2020, 2023), but most of these data were collected in male rats. A central goal of this study was to test the hypothesis that both male and female Avoider rats would show increased alcohol drinking after stress in a free-choice condition. This hypothesis was supported only in male rats. Greater avoidance behavior in male rats also predicted higher consumption of alcohol adulterated with 60 mg/L quinine, a tastant that has been shown to be equally aversive in both male and female rodents (Sneddon et al., 2019; Radke et al., 2020; Katner et al., 2022). This increase in post-stress quinine-adulterated alcohol drinking is interpreted as a form of aversion-resistant or ‘compulsive-like’ alcohol drinking, which is reminiscent of various forms of continued drinking despite negative consequences seen in humans with AUD (see De Oliviera Sergio et al., 2023 for review).

In lab animals, the addition of a moderate dose of quinine to alcohol drinking solutions typically produces a quantifiable reduction in alcohol drinking. This response can be blunted in subgroups of animals by a variety of factors known to support increased alcohol drinking, such as chronic intermittent alcohol access (Hopf et al., 2010), early life stress (Radke et al., 2020, 2021), chronic predatory stress (Shaw et al., 2020), and a genetic predisposition for high alcohol preference (Katner et al., 2022), which suggests that these factors may disrupt the motivational balance between the reinforcing effects of alcohol and the aversive, anti-reinforcing effects of quinine. The present results demonstrate that an acute stressor (predator odor) can also blunt the effectiveness of an aversive stimulus in reducing alcohol intake in individual stress-reactive animals, in this case, in more avoidant male rats. Previous work using an operant procedure found that male Avoider rats will also show operant responding for alcohol that is more resistant to quinine challenge (Edwards et al., 2013).

Five days after the end of alcohol drinking (28 days after stress), we tested rats for their anxiety-like behavior using the EPM. Female Avoider rats showed heightened anxiety-like behavior, as indicated by more stretch attempts (i.e., aborted entries into open areas) compared to unstressed Controls. Male Avoider rats, on the other hand, did not show any differences in behavior in the EPM compared to Non-Avoiders and Controls. Previous work shows that predator odor stress heightens anxiety-like behavior, as measured using the EPM and open field tests, in both male Avoiders and Non-Avoiders up to 5 days post-stress (Whitaker & Gilpin, 2015). By day 9 post-stress, anxiety-like behavior in male and female Non-Avoiders return to Control levels, whereas anxiety-like behavior in Avoiders remain elevated (Weera et al., 2023). Here, we tested anxiety-like behavior at a more protracted time point, that is 28 days after stress, in both male and female rats, and found that the heightened anxiety-like behavior in male but not in female Avoiders had returned to Control levels at this time point. It is currently unclear how long-lasting predator odor stress effects are on anxiety-like behavior in female Avoider rats, but our results show that stress-heightened anxiety-like behavior persist for at least 1 month after stress. It is important to note that the animals in the present study have undergone multiple months of alcohol drinking, similar to those in Weera et al. (2023), but the rats in Whitaker and Gilpin (2015) were alcohol naïve. Another important previous finding to consider is that, in alcohol-drinking male and female rats that had undergone multiple acoustic startle tests, open arms times and entries on the EPM were similar between stressed, and unstressed rats 17 days after predator odor exposure (Albrechet-Souza et al., 2020).

Rats produce a variety of ultrasonic vocalizations (USVs) which may correspond with their affective state (Knutson et al., 2002; Litvin et al., 2007; Brudzynski, 2013). For instance, vocalizations in the 18-32 kHz frequency band are emitted during a variety of aversive experiences and are proposed to serve as alarm calls indicating distress or threat (Brudzynski, 2001, 2021; Burgdorf et al., 2008; Takahashi et al., 2010). We recorded USVs emitted by Avoider, Non-Avoider, and unstressed Control rats before stress (i.e., during place conditioning Pretest), during stress, and after stress during place conditioning testing (Posttest) to track the affective state of these animals throughout the procedure. While the total number of USVs were fairly consistent across Avoiders and Non-Avoiders and throughout the predator odor place conditioning procedure, we found that, in males, Non-Avoiders exhibited a greater proportion of USVs in the <32 kHz range compared to Avoiders during both Pretest (i.e., before stress) and Posttest (i.e., after stress). Avoidance scores also correlated with percent of USVs in the <32 kHz range, such that male rats that emitted more of their vocalizations in the <32 kHz range during Pretest were more likely to not avoid a stress-paired context, and this pattern persisted into Posttest. At this time, it is unclear why male Non-Avoider rats emit more vocalizations within the “aversive” range than male Avoiders. Perhaps, males that respond to novel environments (i.e., the novel place conditioning apparatus during the Pretest) with greater numbers of low frequency alarm calls are more likely to be active coping individuals, thereby exhibiting a Non-Avoider status (or approach coping strategy) following predator odor stress. USVs functioning as alarm calls are often emitted from a place of safety following an adverse exposure (Litvin et al., 2007; Brudzynski, 2021). In the Pretest, Non-Avoiders experienced the anxiogenic novel apparatus but may have perceived relatively more safety than Avoiders and thus initiate alarm call social communication. Female rats emitted far fewer USVs in the <32 kHz range compared to males, regardless of stress group. During fear conditioning, female Wistar rats emit very few low frequency USVs (Schwarting, 2018; Willadsen et al., 2021) which suggests that this type of social communication during aversive experiences is generally specific to males.

To begin to elucidate the neural mechanisms behind the subpopulation-specific changes in alcohol drinking and anxiety-like behaviors after predator odor stress, we quantified gene expression of various CRF neuropeptide system components in the LH. In male rats, Avoiders had significantly increased levels of *crh* in the LH compared to both controls and Non-Avoiders. Given that brains were collected 4 weeks after exposure to predator odor, this prolonged increase in *crh* may represent a potential mechanism underlying avoidance behavior as well as the increase in alcohol drinking observed post-stress in male Avoider rats. Indeed, *crh* was significantly correlated with the change in time spent in the odor-paired chamber (i.e., avoidance scores), such that greater avoidance predicted higher levels of *crh* in the LH. Moreover, *crh* expression also tended to be correlated with consumption of quinine-adulterated alcohol (60 mg/L) and post-stress change in total unadulterated alcohol consumption. In addition, there was a correlation between CRH binding protein gene (*crhbp*) expression in the LH and change in alcohol drinking from pre-stress to post-stress, such that greater increases in drinking post-stress predicted higher levels of *crhbp* in male rats. In Non-Avoider female rats, levels of LH *crh* and *crhbp* predicted lower anxiety-like behavior, as indicated by more open arm entries in the EPM. Collectively, these findings highlight the importance of the CRH system within the LH in mediating subpopulation (i.e., Sex- and Stress Group-specific) differences in stress-alcohol interactions and inform our ongoing circuit-based studies centered on the LH.

Extrahypothalamic CRF, especially in the extended amygdala, is well known to mediate escalated forms of alcohol drinking, and preclinical studies showed that inhibition of CRF1 receptors rescues the escalated alcohol self-administration seen in alcohol-dependent animals (Funk et al., 2006, 2007). However, clinical trials with CRF1 antagonists have not yielded success for alleviating AUD symptoms (Kwako, Spagnolo, et al., 2015; Schwandt et al., 2016), highlighting the need to further elucidate CRF-CRF1 circuit and cellular mechanisms in alcohol drinking and related neuropsychiatric conditions (Pomrenze et al., 2017; Weera & Gilpin, 2024). Indeed, circuit-based studies show that the role of CRF-CRF1 signaling in alcohol drinking may have subpopulation-dependent effects. For instance, silencing CRF+ neurons projecting from the central amygdala (CeA) to the LH or CRF1 antagonism within the LH blunts binge-like alcohol consumption in male but not female mice (Bendrath et al., 2025), while silencing CRF1+ CeA-to-LH neurons reduces alcohol self-administration in stress-exposed male and female Avoider but not Non-Avoider rats (Weera et al., 2023). Our ongoing work is focused on understanding the roles of LH cells and circuits, including those that express CRF/CRF1, in stress-alcohol interactions across subpopulations of animals.

Collectively, these studies show that acute predator odor stress, which recapitulates some aspects of traumatic stress, increases free-choice alcohol drinking, aversion-resistant alcohol drinking, and anxiety-like behavior in a manner that is dependent sex and stress responsivity (i.e., Avoider vs. Non-Avoider phenotype). We highlight the utility of pre-stress ultrasonic vocalizations for predicting Avoider vs. Non-Avoider status after stress. Finally, our gene expression analysis suggests that the CRF system within the lateral hypothalamus plays an important role in post-stress behavioral outcomes.

## Acknowledgements

The authors would like to thank the Comparative Medicine Services (CMS) team at Tufts University in Medford for their reliable and caring husbandry of the animals in this experiment. We would also like to thank Eli Boshak, Mason Rauch, Molly Sikma, and Ayesha Dangyach for their integral hands-on contributions to the long-term alcohol drinking aspects of these experiments. Finally, we are grateful to Vera Liao and Decker Dunlop for their diligent analysis of the USV detection files.

## References

Albrechet-Souza, L., & Gilpin, N. W. (2019). The predator odor avoidance model of post-traumatic stress disorder in rats. Behavioural Pharmacology, 30(2 and 3), 105–114. 10.1097/FBP.0000000000000460

Albrechet-Souza, L., Schratz, C. L., & Gilpin, N. W. (2020). Sex differences in traumatic stress reactivity in rats with and without a history of alcohol drinking. Biology of Sex Differences, 11(1), 27. 10.1186/s13293-020-00303-w

American Psychiatric Association. (2013). Diagnostic and Statistical Manual of Mental Disorders (Fifth Edition). American Psychiatric Association. https://psychiatryonline.org/doi/book/10.1176/appi.books.9780890425596

Bendrath, S. C., Méndez, H. G., Dankert, A. M., Lerma-Cabrera, J. M., Carvajal, F., Dornellas, A. P. S., Lee, S., Neira, S., Haun, H., Delpire, E., Navarro, M., Kash, T. L., & Thiele, T. E. (2025). Corticotropin-Releasing Factor Modulates Binge-Like Ethanol Drinking in a Sex-Dependent Manner: Impact of Amygdala Deletion and Inhibition of a Central Amygdala to Lateral Hypothalamus Circuit. Biological Psychiatry Global Open Science, 5(1), 100405. 10.1016/j.bpsgos.2024.100405

Blanco, C., Xu, Y., Brady, K., Pérez-Fuentes, G., Okuda, M., & Wang, S. (2013). Comorbidity of posttraumatic stress disorder with alcohol dependence among US adults: Results from National Epidemiological Survey on Alcohol and Related Conditions. Drug and Alcohol Dependence, 132(3), 630–638. 10.1016/j.drugalcdep.2013.04.016

Brudzynski, S. M. (2001). Pharmacological and behavioral characteristics of 22 kHz alarm calls in rats. Neuroscience & Biobehavioral Reviews, 25(7–8), 611–617. 10.1016/S0149-7634(01)00058-6

Brudzynski, S. M. (2013). Ethotransmission: Communication of emotional states through ultrasonic vocalization in rats. Current Opinion in Neurobiology, 23(3), 310–317. 10.1016/j.conb.2013.01.014

Brudzynski, S. M. (2021). Biological Functions of Rat Ultrasonic Vocalizations, Arousal Mechanisms, and Call Initiation. Brain Sciences, 11(5), 605. 10.3390/brainsci11050605

Burgdorf, J., Kroes, R. A., Moskal, J. R., Pfaus, J. G., Brudzynski, S. M., & Panksepp, J. (2008). Ultrasonic vocalizations of rats (Rattus norvegicus) during mating, play, and aggression: Behavioral concomitants, relationship to reward, and self-administration of playback. Journal of Comparative Psychology, 122(4), 357.

Coffey, K. R., Marx, R. E., & Neumaier, J. F. (2019). DeepSqueak: A deep learning-based system for detection and analysis of ultrasonic vocalizations. Neuropsychopharmacology, 44(5), 859–868. 10.1038/s41386-018-0303-6

De Oliveira Sergio, T., Frasier, R. M., & Hopf, F. W. (2023). Animal models of compulsion alcohol drinking: Why we love quinine-resistant intake and what we learned from it. Frontiers in Psychiatry, 14, 1116901. 10.3389/fpsyt.2023.1116901

Edwards, S., Baynes, B. B., Carmichael, C. Y., Zamora-Martinez, E. R., Barrus, M., Koob, G. F., & Gilpin, N. W. (2013). Traumatic stress reactivity promotes excessive alcohol drinking and alters the balance of prefrontal cortex-amygdala activity. Translational Psychiatry, 3(8), e296–e296. 10.1038/tp.2013.70

Eghtesad, M., Elahdadi Salmani, M., Lashkarbolouki, T., & Goudarzi, I. (2022). Lateral Hypothalamus Corticotropin-releasing Hormone Receptor-1 Inhibition and Modulating Stress-induced Anxiety Behavior. Basic and Clinical Neuroscience, 13(3), 373–384. 10.32598/bcn.2021.445.3

Erol, A., & Karpyak, V. M. (2015). Sex and gender-related differences in alcohol use and its consequences: Contemporary knowledge and future research considerations. Drug and Alcohol Dependence, 156, 1–13. 10.1016/j.drugalcdep.2015.08.023

Funk, C. K., O’Dell, L. E., Crawford, E. F., & Koob, G. F. (2006). Corticotropin-releasing factor within the central nucleus of the amygdala mediates enhanced ethanol self-administration in withdrawn, ethanol-dependent rats. The Journal of Neuroscience: The Official Journal of the Society for Neuroscience, 26(44), 11324–11332. 10.1523/JNEUROSCI.3096-06.2006

Funk, C. K., Zorrilla, E. P., Lee, M.-J., Rice, K. C., & Koob, G. F. (2007). Corticotropin-releasing factor 1 antagonists selectively reduce ethanol self-administration in ethanol-dependent rats. Biological Psychiatry, 61(1), 78–86. 10.1016/j.biopsych.2006.03.063

Gilpin, N. W., & Weiner, J. L. (2017). Neurobiology of comorbid post-traumatic stress disorder and alcohol-use disorder. *Genes*, Brain and Behavior, 16(1), 15–43. 10.1111/gbb.12349

Goldstein, R. B., Smith, S. M., Chou, S. P., Saha, T. D., Jung, J., Zhang, H., Pickering, R. P., Ruan, W. J., Huang, B., & Grant, B. F. (2016). The epidemiology of DSM-5 posttraumatic stress disorder in the United States: Results from the National Epidemiologic Survey on Alcohol and Related Conditions-III. Social Psychiatry and Psychiatric Epidemiology, 51(8), 1137–1148. 10.1007/s00127-016-1208-5

Guinle, M. I. B., & Sinha, R. (2020). The Role of Stress, Trauma, and Negative Affect in the Development of Alcohol Misuse and Alcohol Use Disorders in Women. Alcohol Research: Current Reviews, 40(2), arcr.v40.2.05. 10.35946/arcr.v40.2.05

Hopf, F. W., Chang, S., Sparta, D. R., Bowers, M. S., & Bonci, A. (2010). Motivation for Alcohol Becomes Resistant to Quinine Adulteration After 3 to 4 Months of Intermittent Alcohol Self-Administration. Alcoholism: Clinical and Experimental Research, 34(9), 1565–1573. 10.1111/j.1530-0277.2010.01241.x

Itoga, C. A., Roltsch Hellard, E. A., Whitaker, A. M., Lu, Y.-L., Schreiber, A. L., Baynes, B. B., Baiamonte, B. A., Richardson, H. N., & Gilpin, N. W. (2016). Traumatic Stress Promotes Hyperalgesia via Corticotropin-Releasing Factor-1 Receptor (CRFR1) Signaling in Central Amygdala. Neuropsychopharmacology, 41(10), 2463–2472. 10.1038/npp.2016.44

Katner, S. N., Sentir, A. M., Steagall, K. B., Ding, Z.-M., Wetherill, L., Hopf, F. W., & Engleman, E. A. (2022). Modeling Aversion Resistant Alcohol Intake in Indiana Alcohol-Preferring (P) Rats. Brain Sciences, 12(8), 1042. 10.3390/brainsci12081042

Knutson, B., Burgdorf, J., & Panksepp, J. (2002). Ultrasonic vocalizations as indices of affective states in rats. Psychological Bulletin, 128(6), 961–977. 10.1037/0033-2909.128.6.961

Koob, G. F. (2013a). Negative reinforcement in drug addiction: The darkness within. Current Opinion in Neurobiology, 23(4), 559–563. 10.1016/j.conb.2013.03.011

Koob, G. F. (2013b). Theoretical frameworks and mechanistic aspects of alcohol addiction: Alcohol addiction as a reward deficit disorder. Current Topics in Behavioral Neurosciences, 13, 3–30. 10.1007/7854_2011_129

Koob, G. F. (2014). Neurocircuitry of alcohol addiction. In Handbook of Clinical Neurology (Vol. 125, pp. 33–54). Elsevier. https://linkinghub.elsevier.com/retrieve/pii/B9780444626196000033

Koob, G. F., & Volkow, N. D. (2010). Neurocircuitry of Addiction. Neuropsychopharmacology, 35(1), 217–238. 10.1038/npp.2009.110

Kwako, L. E., Schwandt, M. L., Sells, J. R., Ramchandani, V. A., Hommer, D. W., George, D. T., Sinha, R., & Heilig, M. (2015). Methods for inducing alcohol craving in individuals with co-morbid alcohol dependence and posttraumatic stress disorder: Behavioral and physiological outcomes. Addiction Biology, 20(4), 733–746. 10.1111/adb.12150

Kwako, L. E., Spagnolo, P. A., Schwandt, M. L., Thorsell, A., George, D. T., Momenan, R., Rio, D. E., Huestis, M., Anizan, S., Concheiro, M., Sinha, R., & Heilig, M. (2015). The Corticotropin Releasing Hormone-1 (CRH1) Receptor Antagonist Pexacerfont in Alcohol Dependence: A Randomized Controlled Experimental Medicine Study. Neuropsychopharmacology, 40(5), 1053–1063. 10.1038/npp.2014.306

Lehavot, K., Stappenbeck, C. A., Luterek, J. A., Kaysen, D., & Simpson, T. L. (2014). Gender differences in relationships among PTSD severity, drinking motives, and alcohol use in a comorbid alcohol dependence and PTSD sample. Psychology of Addictive Behaviors, 28(1), 42–52. 10.1037/a0032266

Litvin, Y., Blanchard, D. C., & Blanchard, R. J. (2007). Rat 22 kHz ultrasonic vocalizations as alarm cries. Behavioural Brain Research, 182(2), 166–172.

Pomrenze, M. B., Fetterly, T. L., Winder, D. G., & Messing, R. O. (2017). The Corticotropin Releasing Factor Receptor 1 in Alcohol Use Disorder: Still a Valid Drug Target? *Alcoholism*, Clinical and Experimental Research, 41(12), 1986–1999. 10.1111/acer.13507

Radke, A. K., Held, I. T., Sneddon, E. A., Riddle, C. A., & Quinn, J. J. (2020). Additive influences of acute early life stress and sex on vulnerability for aversion-resistant alcohol drinking. Addiction Biology, 25(6), e12829. 10.1111/adb.12829

Radke, A. K., Sneddon, E. A., Frasier, R. M., & Hopf, F. W. (2021). Recent Perspectives on Sex Differences in Compulsion-Like and Binge Alcohol Drinking. International Journal of Molecular Sciences, 22(7), 3788. 10.3390/ijms22073788

Ranney, R., Zakeri, S. E., Kevorkian, S., Rappaport, L., Chowdhury, N., Amstadter, A., Dick, D., Spit for Science Working Group, & Berenz, E. C. (2021). Investigating Relationships Among Distress Tolerance, PTSD Symptom Severity, and Alcohol Use. Journal of Psychopathology and Behavioral Assessment, 43(2), 259–270. 10.1007/s10862-020-09842-3

Saraiya, T. C., Back, S. E., Jarnecke, A. M., Blakey, S. M., Bauer, A. G., Brown, D. G., Ruglass, L. M., Killeen, T., & Hien, D. A. (2023). Sex and Gender Differences in Co-Occurring Alcohol Use Disorder and PTSD. Current Addiction Reports, 10(4), 617–627. 10.1007/s40429-023-00511-5

Schreiber, A. L., Lu, Y.-L., Baynes, B. B., Richardson, H. N., & Gilpin, N. W. (2017). Corticotropin-releasing factor in ventromedial prefrontal cortex mediates avoidance of a traumatic stress-paired context. Neuropharmacology, 113, 323–330. 10.1016/j.neuropharm.2016.05.008

Schwandt, M. L., Cortes, C. R., Kwako, L. E., George, D. T., Momenan, R., Sinha, R., Grigoriadis, D. E., Pich, E. M., Leggio, L., & Heilig, M. (2016). The CRF1 Antagonist Verucerfont in Anxious Alcohol-Dependent Women: Translation of Neuroendocrine, But not of Anti-Craving Effects. Neuropsychopharmacology: Official Publication of the American College of Neuropsychopharmacology, 41(12), 2818–2829. 10.1038/npp.2016.61

Schwarting, R. K. W. (2018). Ultrasonic vocalization in female rats: A comparison among three outbred stocks from pups to adults. Physiology & Behavior, 196, 59–66. 10.1016/j.physbeh.2018.08.009

Shaw, G. A., Bent, M. A. M., Council, K. R., Pais, A. C., Amstadter, A., Wolstenholme, J. T., Miles, M. F., & Neigh, G. N. (2020). Chronic repeated predatory stress induces resistance to quinine adulteration of ethanol in male mice. Behavioural Brain Research, 382, 112500. 10.1016/j.bbr.2020.112500

Simms, J. A., Steensland, P., Medina, B., Abernathy, K. E., Chandler, L. J., Wise, R., & Bartlett, S. E. (2008). Intermittent Access to 20% Ethanol Induces High Ethanol Consumption in Long–Evans and Wistar Rats. Alcoholism: Clinical and Experimental Research, 32(10), 1816–1823. 10.1111/j.1530-0277.2008.00753.x

Sinha, R. (2009). Modeling stress and drug craving in the laboratory: Implications for addiction treatment development. Addiction Biology, 14(1), 84–98. 10.1111/j.1369-1600.2008.00134.x

Sneddon, E. A., White, R. D., & Radke, A. K. (2019). Sex Differences in Binge-Like and Aversion-Resistant Alcohol Drinking in C57BL/J6 Mice. Alcoholism: Clinical and Experimental Research, 43(2), 243–249. 10.1111/acer.13923

Straus, E., Haller, M., Lyons, R. C., & Norman, S. B. (2018). Functional and Psychiatric Correlates of Comorbid Post-Traumatic Stress Disorder and Alcohol Use Disorder. Alcohol Research: Current Reviews, 39(2), 121–129.

Takahashi, N., Kashino, M., & Hironaka, N. (2010). Structure of rat ultrasonic vocalizations and its relevance to behavior. PloS One, 5(11), e14115.

Vendruscolo, L. F., Barbier, E., Schlosburg, J. E., Misra, K. K., Whitfield, T. W., Logrip, M. L., Rivier, C., Repunte-Canonigo, V., Zorrilla, E. P., Sanna, P. P., Heilig, M., & Koob, G. F. (2012). Corticosteroid-dependent plasticity mediates compulsive alcohol drinking in rats. The Journal of Neuroscience: The Official Journal of the Society for Neuroscience, 32(22), 7563–7571. 10.1523/JNEUROSCI.0069-12.2012

Weera, M., & Gilpin, N. W. (2024). Central amygdala CRF1 cells control nociception and anxiety-like behavior. Neuropsychopharmacology: Official Publication of the American College of Neuropsychopharmacology, 49(1), 341–342. 10.1038/s41386-023-01693-2

Weera, M., Schreiber, A. L., Avegno, E. M., & Gilpin, N. W. (2020). The role of central amygdala corticotropin-releasing factor in predator odor stress-induced avoidance behavior and escalated alcohol drinking in rats. Neuropharmacology, 166, 107979. 10.1016/j.neuropharm.2020.107979

Weera, M., Shackett, R. S., Kramer, H. M., Middleton, J. W., & Gilpin, N. W. (2021). Central Amygdala Projections to Lateral Hypothalamus Mediate Avoidance Behavior in Rats. The Journal of Neuroscience, 41(1), 61–72. 10.1523/JNEUROSCI.0236-20.2020

Weera, M., Webster, D. A., Shackett, R. S., Benvenuti, F., Middleton, J. W., & Gilpin, N. W. (2023). Traumatic Stress-Induced Increases in Anxiety-like Behavior and Alcohol Self-Administration Are Mediated by Central Amygdala CRF1 Neurons That Project to the Lateral Hypothalamus. The Journal of Neuroscience, 43(50), 8690–8699. 10.1523/JNEUROSCI.1414-23.2023

Whitaker, A. M., & Gilpin, N. W. (2015). Blunted hypothalamo-pituitary adrenal axis response to predator odor predicts high stress reactivity. Physiology & Behavior, 147, 16–22. 10.1016/j.physbeh.2015.03.033

Willadsen, M., Uengoer, M., Schwarting, R. K. W., Homberg, J. R., & Wöhr, M. (2021). Reduced emission of alarm 22-kHz ultrasonic vocalizations during fear conditioning in rats lacking the serotonin transporter. Progress in Neuro-Psychopharmacology and Biological Psychiatry, 108, 110072. 10.1016/j.pnpbp.2020.110072

