## Supplemental Figures for "Predator odor stress produces sex- and subpopulation-specific increases in alcohol drinking, anxiety-like behavior, and lateral hypothalamic *crh* expression"

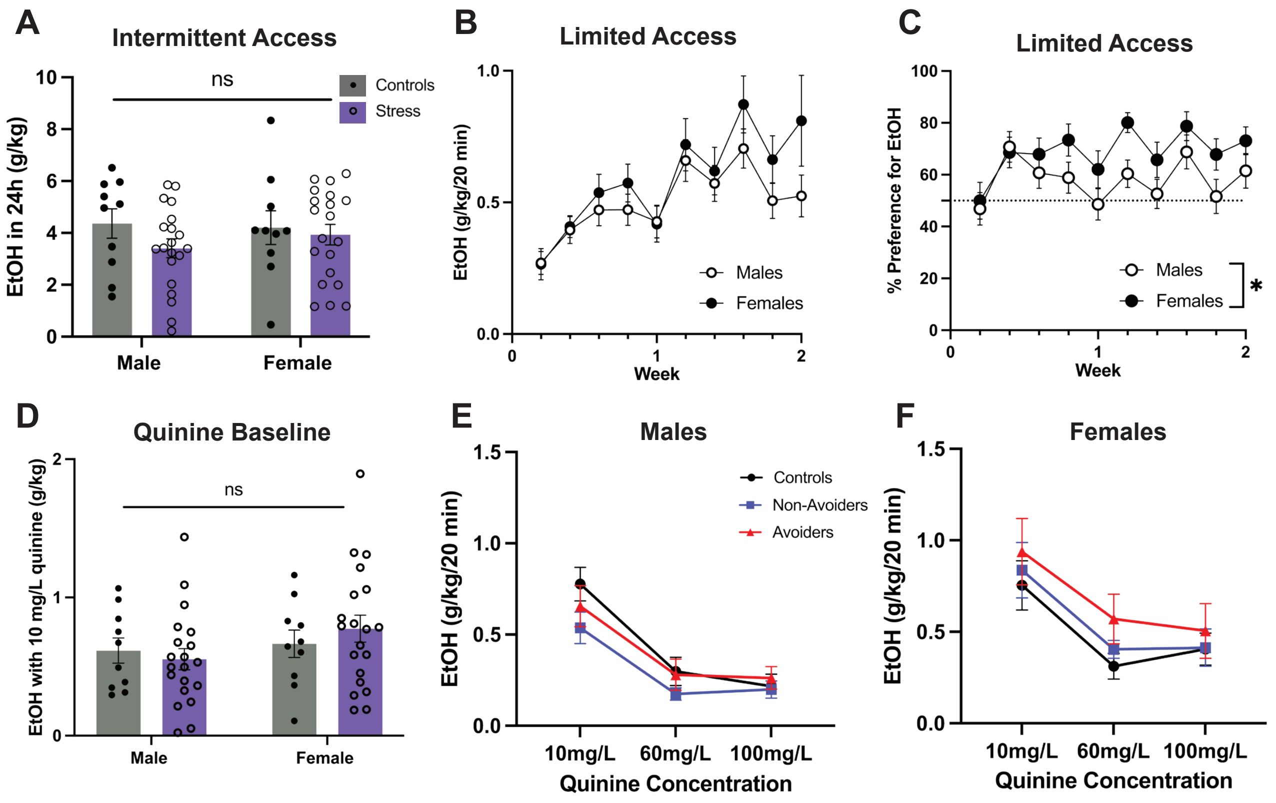


**Supplemental Figure 1. Prestress baselines and post-stress within-subjects comparisons**
**A)** Pre-stress baseline alcohol consumption during the Intermittent Access phase (see Figure 1A). **B)** Pre-stress alcohol consumption in the Limited Access phase. **C)** Pre-stress alcohol preference during the Limited Access phase. **D)** Pre-stress baseline consumption of quinine (10 mg/L) adulterated alcohol (g/kg). **E)** Male and **F)** Female within-subjects quinine drinking across all three post-stress quinine drinking weeks. Error bars represent Mean ± SEM. *$p < 0.05$.

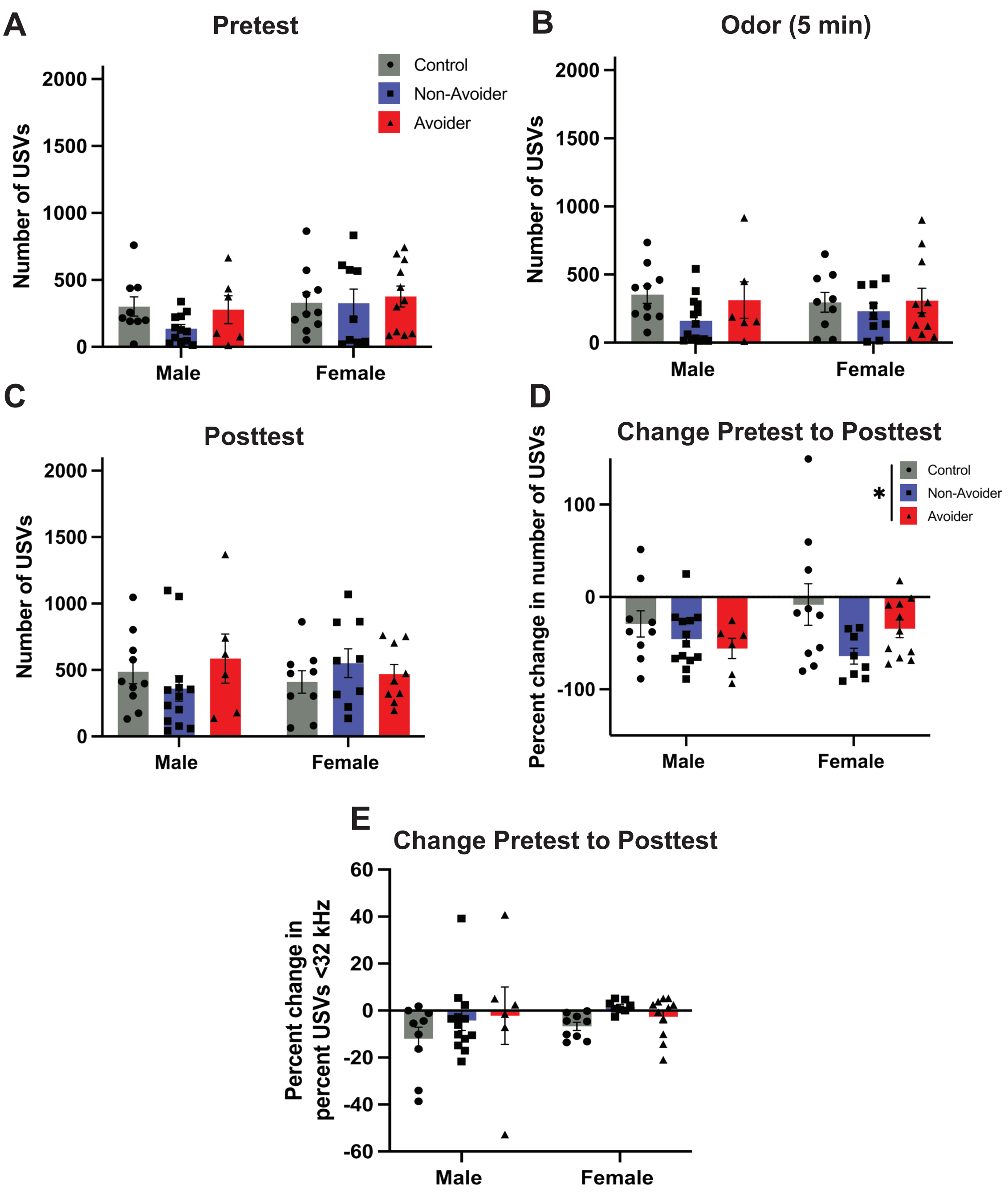


**Supplemental Figure 2: USVs**

**A)** Number of USVs recorded in the Pretest, **B)** first 5 minutes of odor exposure, and **C)** Posttest. **D)** Percent change in the number of USVs emitted in the Posttest compared to the Pretest (Stress Group x Sex ANOVA, main effect of Stress Group $F(2, 51)=3.808$, $p=0.0287$. **E)** Percent change in percentage of USVs <32 kHz in for each individual in the Posttest compared to the Pretest. Error bars represent Mean +/- SEM. $*p < 0.05$.

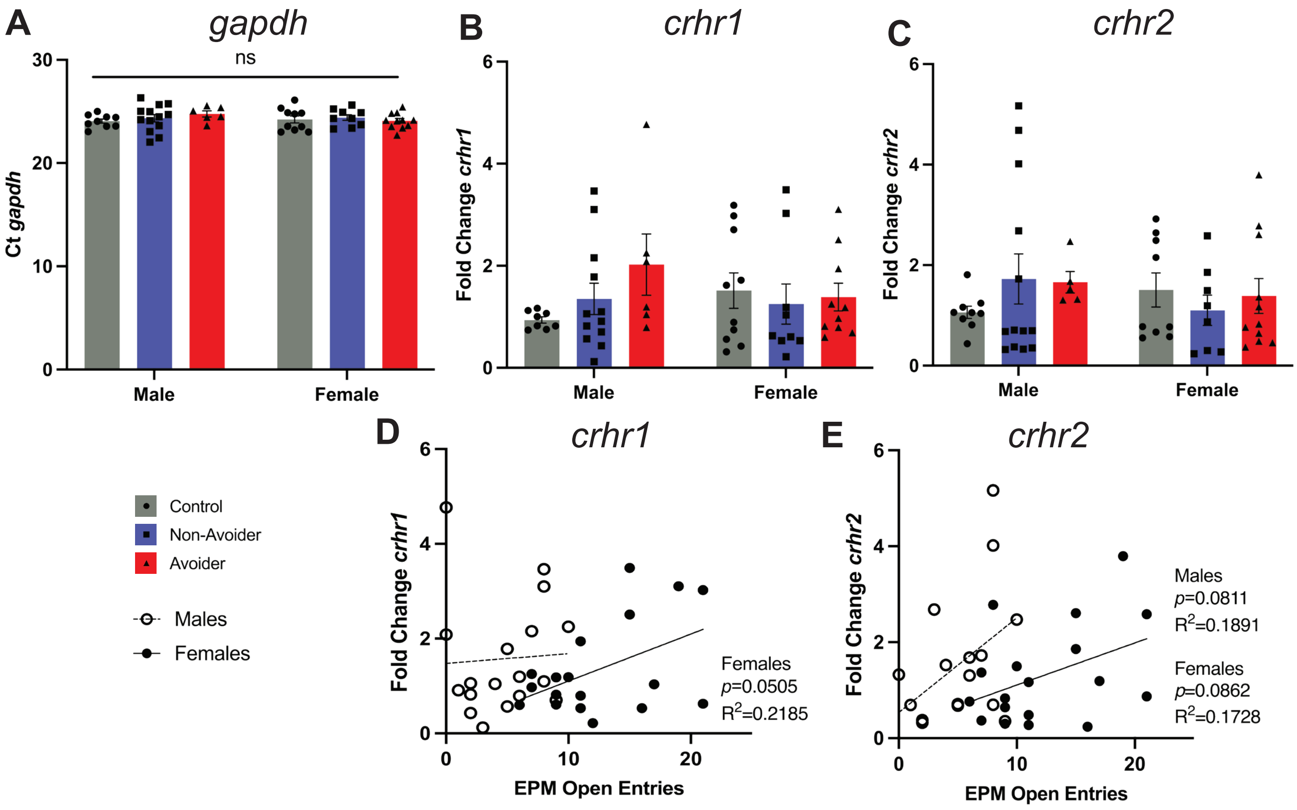


**Supplemental Figure 3: Additional gene expression metrics for *crhr1* and *crhr2***
**A)** Expression (Ct) of housekeeping gene *gapdh*. **B)** Fold change in mRNA expression of *crhr1* and **C)** *crhr2*. **D)** Relationship between open arm entries in the EPM and *crhr1* expression or **E)** *crhr2*. Error bars represent Mean +/- SEM. *$p < 0.05$.
